# Application of an ecology-based analytic approach to discriminate signal and noise in low-biomass microbiome studies: whole lung tissue is the preferred sampling method for amplicon-based characterization of murine lung microbiota

**DOI:** 10.1101/2020.09.08.283259

**Authors:** Jennifer M. Baker, Kevin J. Hinkle, Roderick A. McDonald, Nicole R. Falkowski, Gary B. Huffnagle, Robert P. Dickson

## Abstract

**Background:** Low-biomass microbiome studies (such as those of the lungs, placenta, and skin) are vulnerable to contamination and sequencing stochasticity, which obscure legitimate microbial signal. Since low-biomass microbiome fields have had variable success in establishing the reality and clinical significance of identified microbiota, we sought to develop and apply an analytical approach to discriminate signal from noise in low-biomass microbiome studies. We used this approach to determine the optimal sampling strategy in murine lung microbiome studies, which will be essential for future mechanistic lung microbiome research.

**Methods:** Using a novel, ecology-based analytic approach, we compared bacterial DNA from the lungs of healthy adult mice collected via two common sampling approaches: homogenized whole lung tissue and bronchoalveolar lavage (BAL) fluid. We quantified bacterial DNA using droplet digital PCR, characterized bacterial communities using 16S rRNA gene sequencing, and systematically assessed the quantity and identity of bacterial DNA in both specimen types. We compared bacteria detected in lung specimens to each other and to potential source communities: negative (background) control specimens and paired oral samples.

**Findings:** By all measures, whole lung tissue in mice contained greater bacterial signal and less evidence of contamination than did BAL fluid. Relative to BAL fluid, whole lung tissue exhibited a greater quantity of bacterial DNA, distinct community composition, decreased sample-to-sample variation, and greater biological plausibility when compared to potential source communities. In contrast, bacteria detected in BAL fluid were minimally different from those of procedural, reagent, and sequencing controls.

**Interpretation:** An ecology-based analytical approach discriminates signal from noise in low-biomass microbiome studies and identifies whole lung tissue as the preferred specimen type for murine lung microbiome studies. Sequencing, analysis, and reporting of potential source communities, including negative control specimens and contiguous biological sites, is crucial for biological interpretation of low-biomass microbiome studies, independent of specimen type.

**Funding:** National Institutes of Health

## Introduction

Though the development of next-generation sequencing has led to heightened interest in the study of microbial communities across biological contexts, the study of low-biomass microbiomes is particularly challenging and requires the development of new methodological approaches. Low-biomass samples -samples with low densities of bacterial cells and therefore low quantities of bacterial DNA - are susceptible to contamination with background-derived signal, which affects the taxonomic composition of low-biomass samples^1,2^ and makes it challenging to decipher biological meaning from sequencing data^3^. These methodological challenges exist in all fields that study low-biomass microbial communities across environmental, industrial, and biomedical contexts.

Low-biomass microbiome fields have had variable success in overcoming these methodological challenges. Whereas early findings related to the purported placenta microbiome have subsequently been attributed to contamination^4,5^, the lung microbiome field has flourished with robust, validated findings: lung microbiota are detectable in health^6–12^, correlated with lung immunity both in health^7,8^ and disease^13–15^, correlated with disease severity and predictive of response to therapy^16–19^, and prognostic of clinical outcome in multiple conditions^20–27^. The lung microbiome field addressed the challenge of low-biomass microbiome sampling by systematically defining methods that collect representative populations of lung microbiota to maximize bacterial DNA content and minimize vulnerability to background contamination^9–12^. As a result, empirically validated sampling approaches such as bronchoalveolar lavage (BAL) fluid, which samples a large surface area and yields high sample volumes, and sputum, which contains concentrated densities of bacterial cells, have been successfully implemented in human lung microbiome studies^28^.

Yet despite their routine use in human lung microbiome studies, these sampling methods are not easily adapted for sampling lung microbiota in murine models, which will be critical to understand the mechanisms that govern the relationship between respiratory tract microbiota and pulmonary disease. Anatomic considerations make the application of sequencing-based techniques to murine lung microbiome studies particularly challenging. Collection of BAL fluid is severely limited by the small (∼1 mL) volume of the murine lung^29^, and sputum collection is not possible in mice. In contrast, analysis of homogenized lung tissue is more feasible in mice than humans, and represents a viable option for maximizing the bacterial DNA content in murine lung samples^30^. The ability to effectively sample low-biomass microbial communities is inherently context-dependent and will require new solutions adapted to the particular context of each study.

We therefore designed an empirical approach to compare microbial signal detected in two distinct sample types collected from the same ecological site (murine lungs) with the following goals: 1) to assess the usefulness of microbial ecology-based analytical techniques repurposed for the discrimination of legitimate microbial signal from background noise and 2) to determine the sampling method that is best suited for the characterization of the murine lung microbiome. To accomplish these goals, we quantified and sequenced the bacterial DNA present in BAL fluid and whole lung tissue from otherwise genetically- and environmentally-identical healthy mice and compared them using a novel analytic approach (Figure 1).

**Figure 1:**
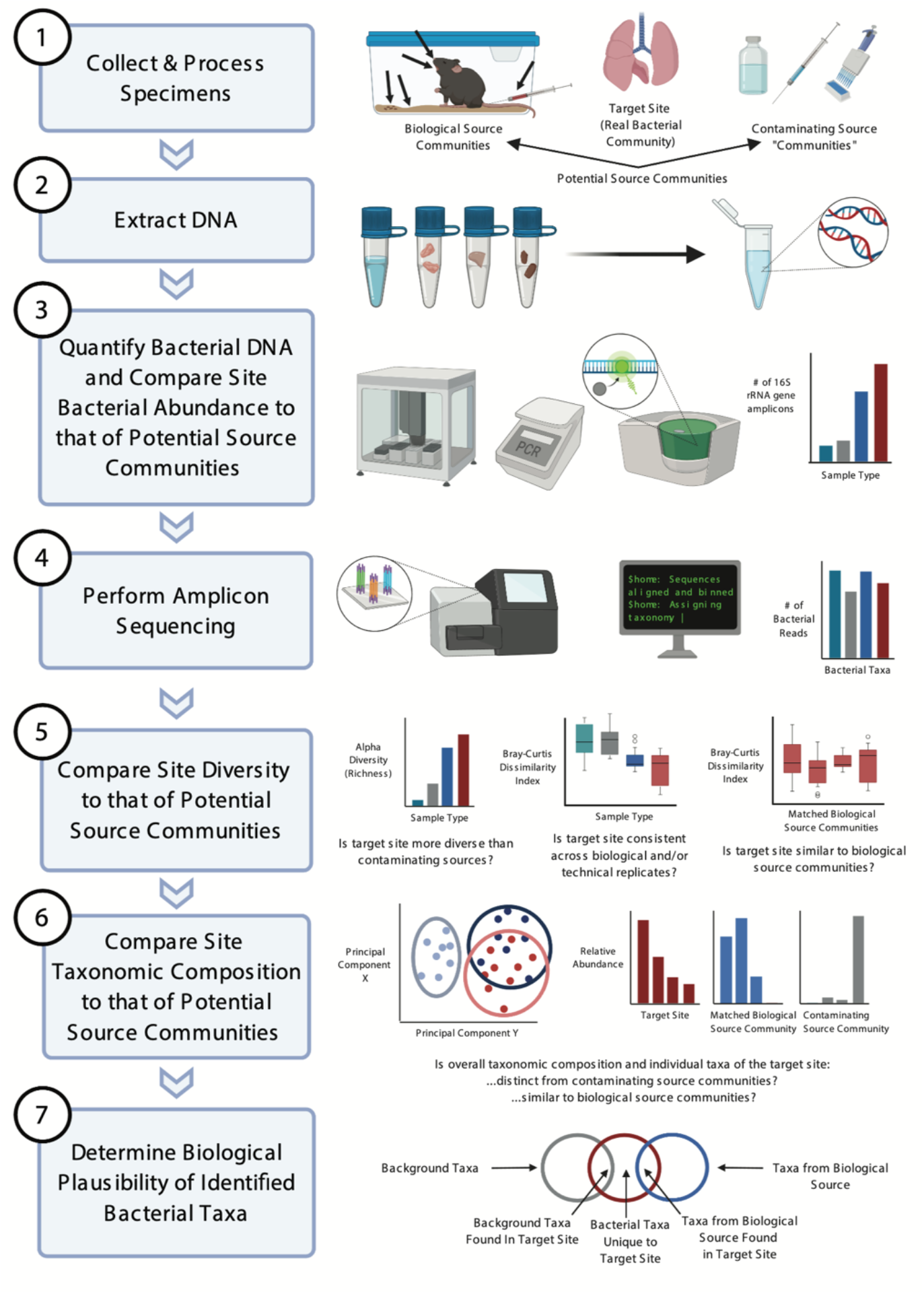
A newly defined experimental and analytic approach can distinguish bacterial signal from noise in low-biomass microbiome studies. Graphical and conceptual outline of an experimental and analytic approach to low-biomass microbiome studies. This approach was applied to murine lung microbiome sampling optimization as a proof-of-concept in this study, and may be useful in other low-biomass microbiome studies across biological contexts.

## Methods

### Ethics approval

The animal studies described in this manuscript were approved by the Institutional Animal Care and Use Committee at the University of Michigan. Laboratory animal care policies at the University of Michigan follow the Public Health Service Policy on Humane Care and Use of Laboratory Animals.

### Mice

Eight-week-old female C57BL/6 mice (n = 20) were purchased from Jackson Laboratories and housed under specific pathogen–free conditions. Mice were housed in five-animal cages in a common animal housing room and did not receive independent ventilation. Mice were allowed to acclimate for 1 week before harvest at 9 weeks of age. To avoid batch effect, mice were randomly assigned to specimen type (BAL fluid or whole lung tissue) and evenly sampled across cages. Animal experimentation was performed in compliance with the ARRIVE Guidelines^31,32^. Details regarding tissue collection and processing are reported in the online supplement (Figure 1, step 1).

### DNA extraction, quantification, and 16S rRNA gene sequencing

DNA was extracted, amplified, and sequenced according to previously published protocols^33,34^ (Figure 1, steps 2,4). Sequencing was performed with the MiSeq platform (Illumina). Bacterial DNA in lung specimens and negative controls was quantified with a QX200 ddPCR system (Bio-Rad, Hercules, CA) according to a previously published protocol^35^. Details are provided in the online supplement.

### Data analysis

16S rRNA gene sequencing data were processed using mothur (v. 1.43.0) according to the Standard Operating Procedure for MiSeq sequence data using a minimum sequence length of 250 base pairs^36,37^. Overall significance was determined as appropriate by the Kruskal-Wallis test and by permutational multivariate ANOVA (PERMANOVA) with 10,000 permutations using Euclidean distances (adonis). Pairwise significance was determined as appropriate by the Wilcoxon test with the Benjamini-Hochberg correction for multiple comparisons, Tukey’s HSD test, and two-sample independent Mann-Whitney U test. All statistical tests used p=0·05 as a threshold for significance. Details regarding statistical and ecologic analysis are reported in the online supplement.

## Results

### Murine whole lung tissue contains more bacterial DNA than BAL fluid and negative controls

Obtaining quality sequencing data depends on the presence of sufficient bacterial DNA in the samples to be analyzed. Therefore, we first compared the quantity of bacterial DNA in whole lung tissue and BAL fluid obtained from healthy C57BL/6 mice (Figure 1, step 3). We hypothesized that whole lung tissue contains more bacterial DNA compared to BAL fluid. To test this hypothesis, we determined the number of 16S rRNA gene copies present in DNA isolated from whole lung tissue, BAL fluid, and negative control specimens using droplet digital PCR (ddPCR). As seen in Figure 2, BAL fluid and whole lung tissue both contained a significantly greater quantity of bacterial DNA than the isolation control (p=0·0084 and 0·0026, respectively). In contrast, BAL fluid did not contain more bacterial DNA than sampling controls or no template controls (p>0·05). Whole lung tissue contained significantly more bacterial DNA than all other groups, including all negative controls (p=0·0001). Whole lung tissue contained 27-fold more 16S rRNA gene copies than BAL fluid (64,110 vs. 2,367 mean copies/mL, respectively; p=0·0002). We thus concluded that murine whole lung tissue contains a greater quantity of bacterial DNA than does BAL fluid.

**Figure 2:**
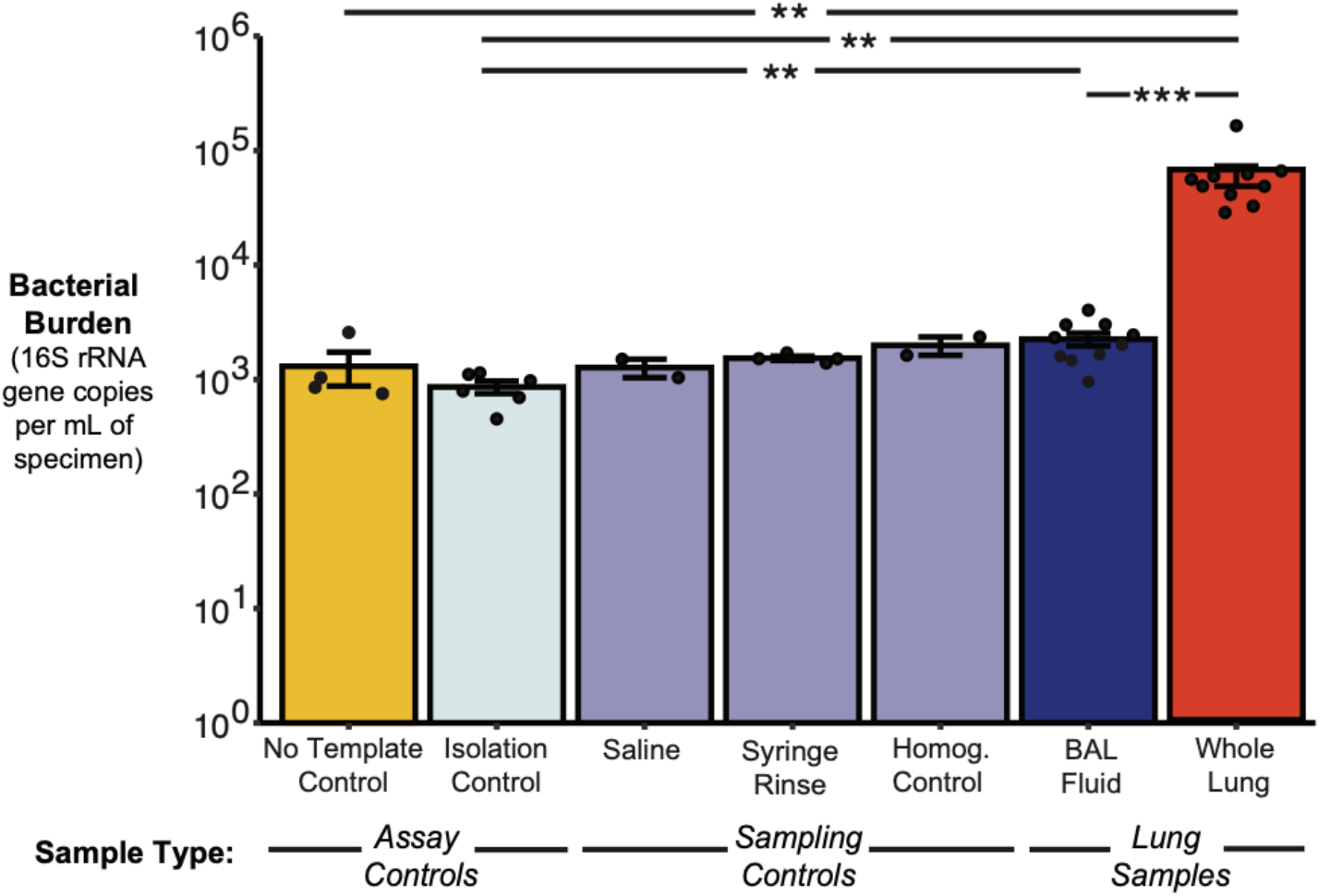
Murine whole lung tissue contains increased bacterial burden relative to BAL fluid and negative controls. Whole lung tissue contains more copies of the bacterial 16S rRNA gene per mL of DNA isolated from lung or control specimens as quantified by ddPCR. Mean ± SEM and individual data points (representing the average of technical duplicates) are shown. Overall significance was determined by the Kruskal-Wallis test (p = 0.0001). Pairwise significance was determined by the pairwise Wilcoxon test and corrected for multiple comparisons using the Benjamini-Hochberg method (pairwise comparisons including whole lung or BAL fluid that are not shown were not significant). Significance key: ns p > 0.05; * p ≤ 0.05; ** p ≤ 0.01; *** p ≤ 0.001; ****p ≤ 0.0001.

Having confirmed the presence of detectable bacterial DNA in whole lung tissue and BAL fluid, we proceeded with 16S rRNA gene sequencing according to a standard low-biomass protocol. Along with whole lung tissue and BAL fluid, we sequenced a variety of controls, including cecum as a high-biomass positive control, tongue as a low-biomass positive control and potential source community of the lower respiratory tract, a synthetic mock community as a positive sequencing control, and negative controls for each stage of specimen processing, including sampling, DNA isolation, and sequencing controls. Details regarding adequacy of sequencing are provided in the online supplement.

### Murine whole lung tissue has increased alpha diversity and decreased sample-to-sample variation relative to BAL fluid and negative controls

We next determined if the alpha (within-sample) diversity also differed across sampling approaches (Figure 1, step 5). We hypothesized that the increased quantity of bacterial DNA in whole lung tissue would yield greater diversity of bacterial taxa in whole lung tissue compared to BAL fluid. To test this hypothesis, we calculated community richness as measured by the number of unique operational taxonomic units (OTUs) present in each specimen and negative control. As predicted, whole lung tissue had greater community richness than BAL fluid (p=0·001) and sampling, isolation, and sequencing controls (p<0·001 for all comparisons) (Figure 3). In contrast, whole lung and BAL specimens did not significantly differ in Shannon diversity index, which reflects both community richness and evenness (p>0·05; Supplementary Figure 2). We therefore concluded that alpha diversity differs across sampling approaches, with greater alpha diversity in whole lung tissue driven by the detection of greater numbers of unique OTUs relative to BAL fluid.

**Figure 3:**
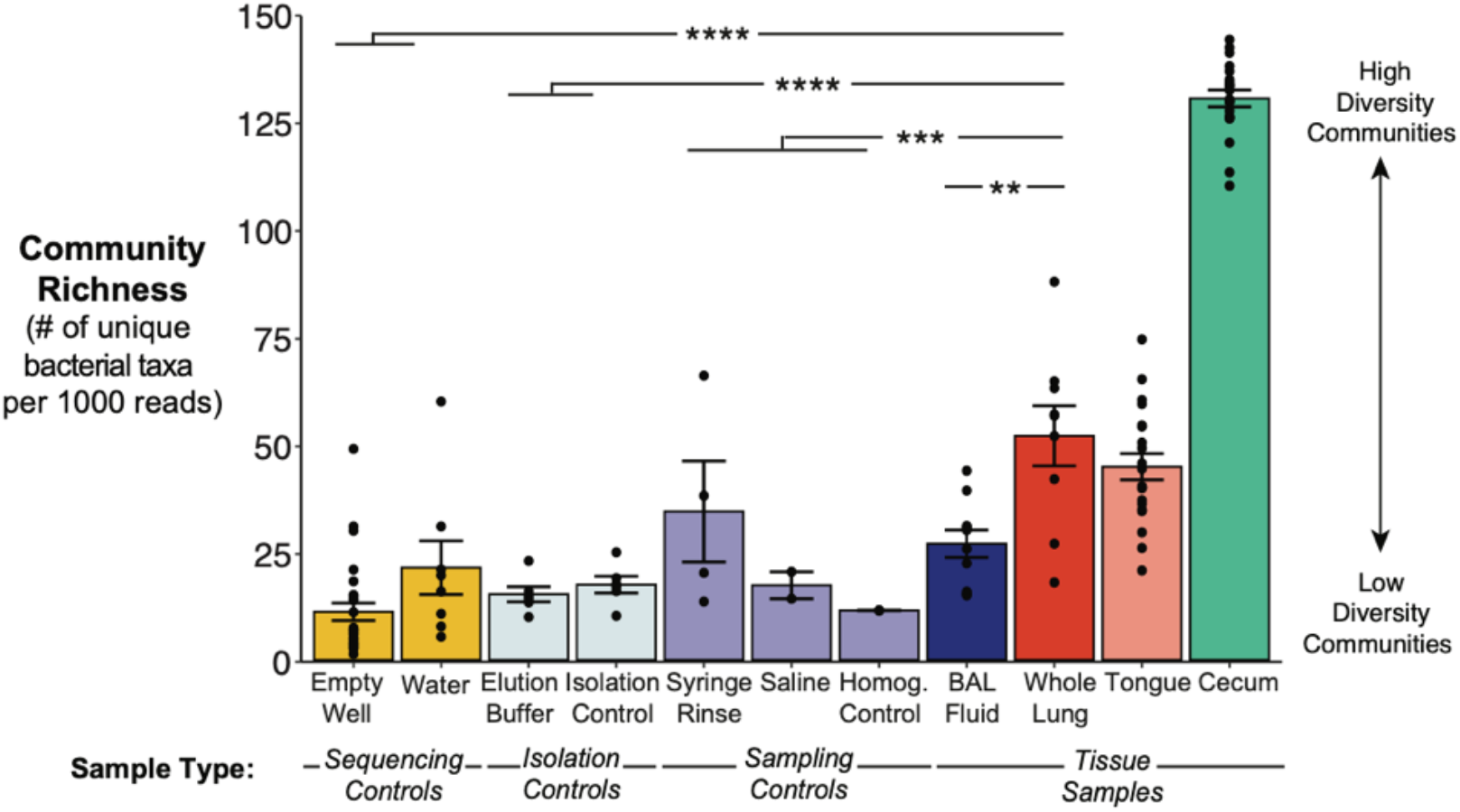
Bacterial communities in murine whole lung tissue have increased alpha diversity relative to BAL fluid and negative controls. A. Whole lung tissue contains a greater number of unique bacterial taxa than BAL fluid and negative controls. Richness of the bacterial community in each tissue or control specimen was determined by clustering reads with species-level similarity (≥ 97% sequence identity) into operational taxonomic units (OTUs) and calculating the number of unique OTUs within each specimen, normalized to 1000 reads per specimen. Mean ± SEM and individual data points are shown. Pairwise significance was determined by comparing whole lung tissue and BAL fluid to pooled sampling, isolation, and sequencing controls (respectively, as shown) using Tukey’s HSD test. Significance key: ns p > 0.05; * p ≤ 0.05; ** p ≤ 0.01; *** p ≤ 0.001; ****p ≤ 0.0001.

Since BAL fluid contained low quantities of bacterial DNA and fewer unique OTUs than whole lung tissue, we suspected that incomplete sampling of the respiratory tract via saline lavage may also result in increased sampling and sequencing stochasticity^38^, which both lead to decreased specimen-to-specimen reproducibility of cohoused mice (which have similar lung microbiota^6^). We thus hypothesized that whole lung tissue would have decreased sample-to-sample variation relative to BAL fluid, representing greater replicability. To test this hypothesis, we computed the Bray-Curtis dissimilarity index, a beta-diversity metric based on pairwise inter-sample distances between specimens of the same type (i.e. we compared each whole lung tissue specimen to each other whole lung tissue specimen, and likewise for BAL fluid). Whole lung tissue yielded a decrease in average Bray-Curtis dissimilarity index relative to that of BAL fluid and empty well controls (p<0.0001) (Figure 4). In contrast, the average Bray-Curtis dissimilarity index for BAL fluid was not significantly different than the highly dissimilar empty well controls (p=0·27). These results indicate that whole lung tissue displays decreased sample-to-sample variation and samples the lung microbiome of mice more reproducibly than BAL fluid.

**Figure 4:**
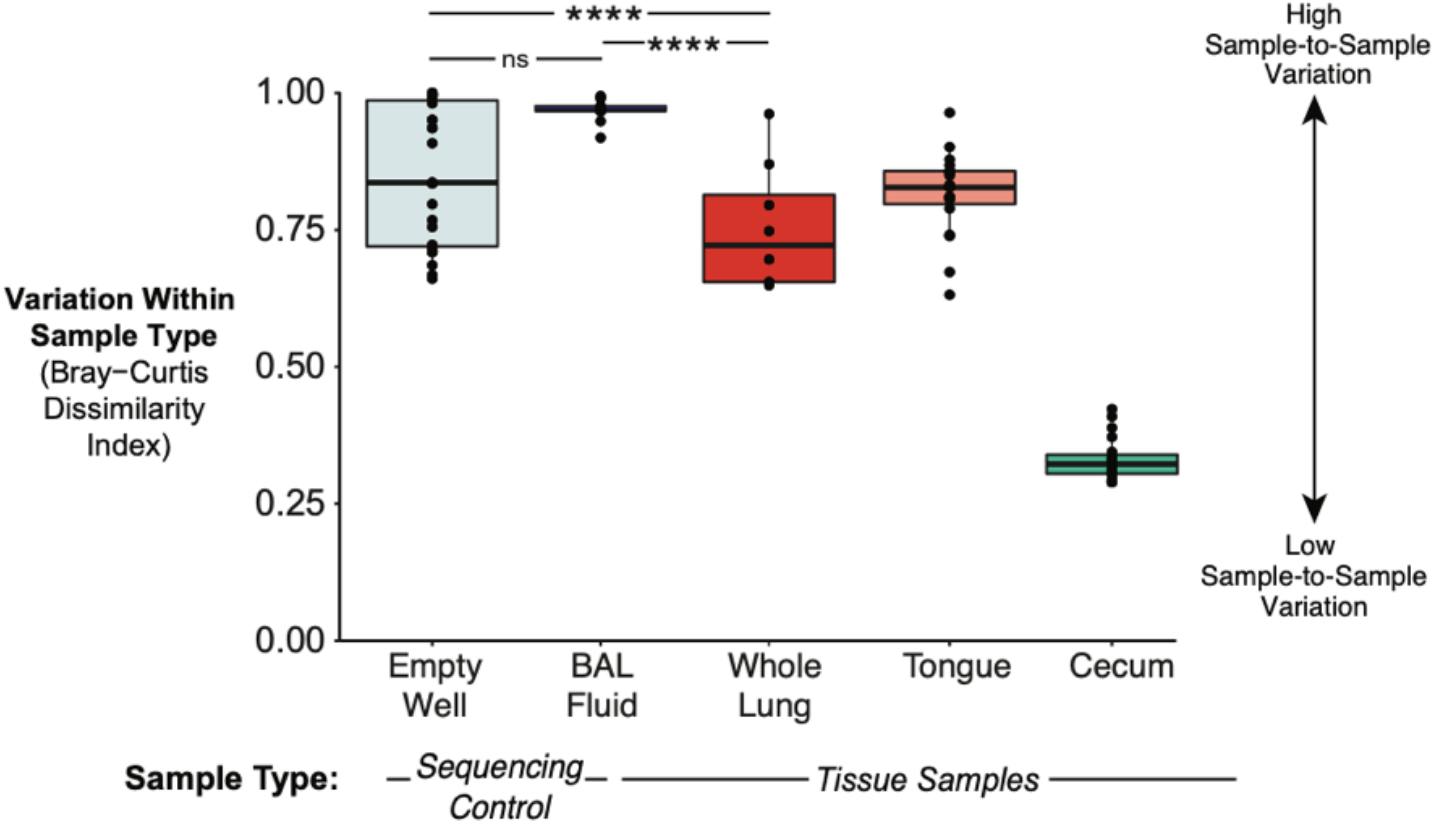
Bacterial communities in murine whole lung tissue show decreased variation among biological replicates compared to those in BAL fluid. Variation among lung bacterial communities of healthy mice from the same shipment was quantified using the Bray-Curtis dissimilarity index. For comparison, Bray-Curtis dissimilarity was also calculated for empty wells as a representative negative control with high variation, cecal communities as a representative body site with low variation, and tongue as a representative seed community for the lower respiratory tract. Median, IQR, and all unique pairwise comparisons (individual data points) are shown. Pairwise significance was determined by pairwise Wilcoxon test and corrected for multiple comparisons using the Benjamini-Hochberg method. Significance key: ns p > 0.05; * p ≤ 0.05; ** p ≤ 0.01; *** p ≤ 0.001; ****p ≤ 0.0001.

### The taxonomic composition of murine whole lung tissue is similar to its oral microbiome source community and is distinct from negative controls, whereas that of BAL fluid is not distinct from negative controls

Having identified differences in bacterial quantity and diversity across sampling approaches, we next assessed whether the taxonomic composition of whole lung tissue and BAL fluid differed from each other and from negative controls (Figure 1, step 6). Since whole lung tissue had higher bacterial DNA content and alpha diversity than BAL fluid, we hypothesized that the taxonomic composition of BAL fluid would more closely resemble that of negative control specimens than would whole lung tissue, reflecting background contamination and sequencing noise as predominant sources of taxa in BAL fluid. To test this hypothesis, we used principal component analysis (PCA) to compare the similarity of taxa identified in whole lung tissue, BAL fluid, and negative control specimens. As seen in Figure 5A, the taxonomic composition of whole lung tissue was distinct from that of BAL fluid (p=0·00009) and pooled sampling controls (p=0·0004). In contrast, BAL fluid showed prominent overlap with sampling controls and did not differ in overall community composition (p=0·46). Similar results were obtained when comparing whole lung tissue and BAL fluid with isolation and sequencing controls (Supplementary Figure 3A,B). Overall, these data show that the taxonomic composition of whole lung tissue is distinct from that of BAL fluid and negative controls, whereas BAL fluid is not distinct from most negative controls.

**Figure 5:**
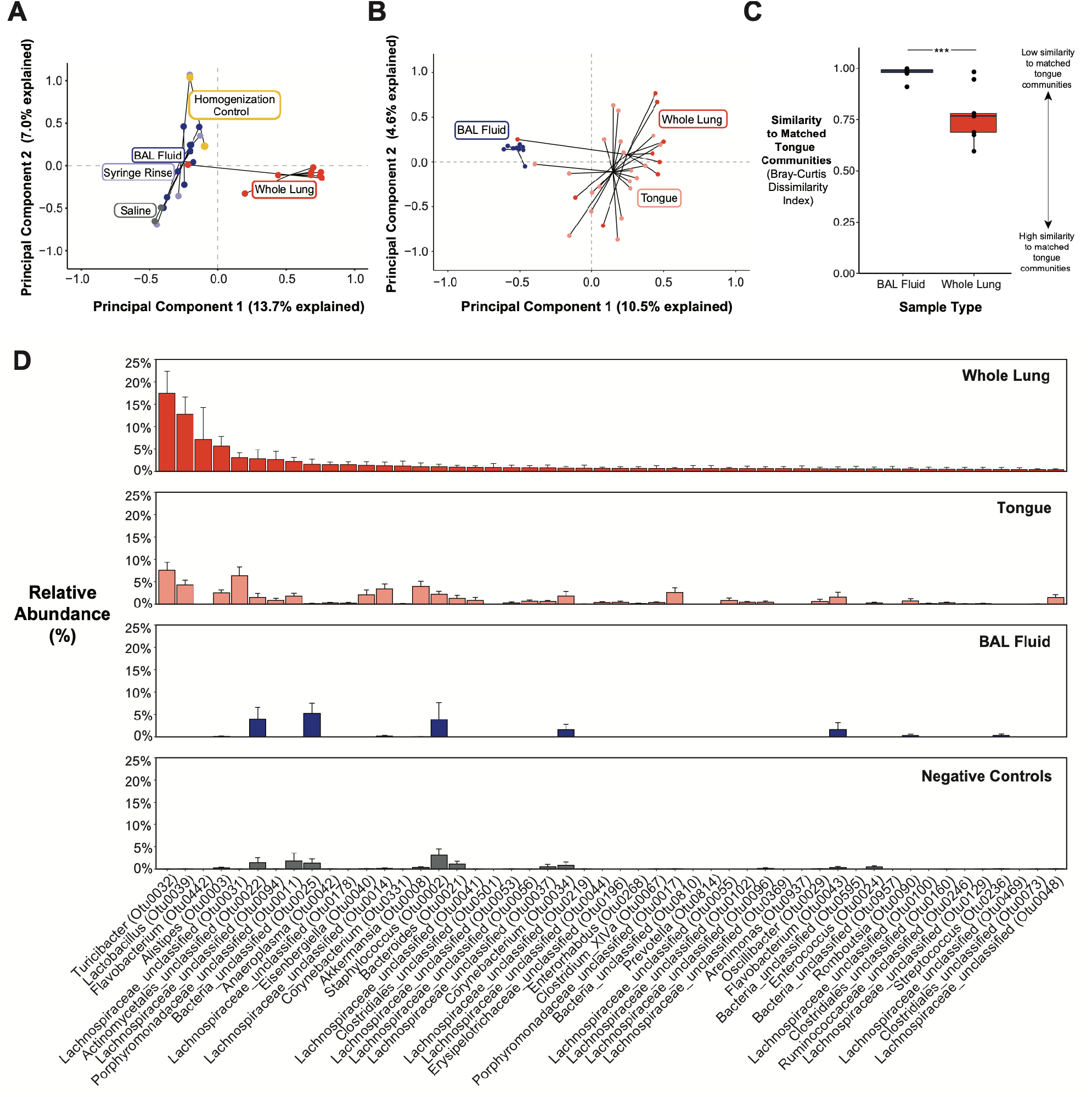
The taxonomic composition of bacterial communities in murine whole lung tissue is distinct from the background-dominant taxonomic composition of BAL fluid and similar to that of the oral microbiome, a biologically plausible source community. A. Whole lung tissue clusters separately from BAL fluid and sampling controls by principal component analysis of Hellinger-transformed 16S rRNA gene sequencing data. Individual data points represent specimens grouped by sample or control type. B. Whole lung tissue, but not BAL fluid, clusters near tongue samples by principal component analysis of Hellinger-transformed 16S rRNA gene sequencing data. Individual data points represent specimens grouped by sample type. C. Bacterial communities in whole lung tissue are more similar to matched (within-mouse) oral communities than BAL fluid. Similarity of lung bacterial communities, grouped by sampling approach, to matched oral communities was quantified using Bray-Curtis dissimilarity index. Median, IQR, and individual data points representing within-mouse comparisons of oral and lung communities are shown. D. Relative abundance of bacterial taxa in whole lung tissue are similar to that of oral bacterial communities. In contrast, the relative abundance of bacterial taxa in BAL fluid are similar to that of negative controls. Bars are ranked by mean abundance in whole lung tissue and represent mean ± SEM percent relative abundance of the top 50 bacterial taxa (OTUs) in whole lung tissue across sample types. Labels denote genus (or most specific taxonomic level if no genus was assigned) and unique identifier for each OTU. Overall significance was determined by (A, B) permutational multivariate ANOVA (p = 0.00009 for both). Pairwise significance was determined by (A, B) two-sample PERMANOVA (A only: pooled sampling controls were compared to each lung sample type), and (C) two-sample unpaired Mann-Whitney U test. Significance key: ns p > 0.05; * p ≤ 0.05; ** p ≤ 0.01; *** p ≤ 0.001; ****p ≤ 0.0001.

We next assessed the biological plausibility of bacterial taxa by comparing whole lung tissue and BAL fluid communities to their likely source community, the oral microbiome (Figure 1, step 7). We hypothesized that the taxonomic composition of whole lung tissue would more closely resemble that of the oral microbiome source community than does BAL fluid. Principal component analysis confirmed that tongue and whole lung tissue display similar but statistically different (p=0·01) taxonomic compositions, whereas BAL fluid clusters separately both from tongue (p=0·00009) and whole lung tissue (Figure 5B). The clustering of BAL fluid with negative controls and tongue with whole lung tissue is also observed when plotting all lung, tongue, and negative control samples together (Supplementary Figure 3C). We confirmed these results by calculating the Bray-Curtis dissimilarity index for matched (i.e. from the same mouse) tongue and lung samples (Figure 5C). Consistent with the PCA results, whole lung tissue more closely resembled the oral source community than did BAL fluid (p=0·0004). Rank abundance analysis revealed that the prominent taxa in whole lung tissue were also common in tongue specimens, whereas taxa in BAL fluid bore little resemblance to oral taxa and instead resembled taxa in negative controls (Figure 5D). The similarity of taxa in the whole lung and tongue samples and the BAL fluid and negative control samples, respectively, can also be observed when ordering rank abundance plots by the taxa found in the tongue or pooled negative controls (Supplementary Figure 4). Together, these results confirm that the bacterial taxa identified in whole lung tissue are more biologically plausible than those detected in BAL fluid.

## Discussion

This study illustrates how an ecology-based analytical approach can determine the reality of bacterial signal in low-biomass microbiome studies. Our approach revealed the superiority of murine whole lung tissue relative to BAL fluid in detecting bacterial signal, and validates the use of whole lung tissue for lung microbiome studies in mice. The bacterial signal in murine whole lung tissue is stronger than that of BAL fluid by all comparisons: increased quantity of bacterial DNA, greater diversity of bacterial taxa, and taxonomic composition that is reproducible across biological replicates, distinct from negative controls, and more similar to the oral microbiome, a biologically plausible source community (Table 1).

**Table 1:** Comparison of Sampling Methods for Murine Lung Microbiome Studies.

|  | **Whole Lung Tissue** | **Bronchoalveolar Lavage Fluid** |
| --- | --- | --- |
| Sample description | All lung lobes homogenized in sterile water | Dislodged airway and alveolar contents (microbes, leukocytes, epithelial cells) in sterile saline |
| Biological site sampled | Airway and intra-alveolar space, interstitium, & blood (if not perfused) | Airway and intra-alveolar space only |
| Bacterial biomass | Low | Low |
| Host-to-microbe DNA ratio | High | Low |
| Total DNA content | High | Low |
| Number of 16S rRNA gene copies | $\sim 10^4$ | $\sim 10^3$ |
| Variation among biological replicates | Low | High |
| Similarity to contaminating source "communities" (negative controls) | Low | High |
| Similarity to biological source community (oral microbiome) | High | Low |

This study represents the first systematic comparison of sampling methods appropriate for the study of the murine lung microbiome. The lack of empirically-validated methods for sampling lung microbiota in mice is particularly concerning in light of the current reproducibility crisis^39^ and recent controversial low-biomass studies^4,5,40^, which highlight the dangers of over-interpreting noisy sequencing data in the absence of rigorous, field-specific standards. A systematic examination of methods for sampling lung microbiota in mice is overdue, especially considering the first report describing the murine lung microbiome was published almost a decade ago^41^. Published murine lung microbiome studies to date have used both whole lung tissue^6,42–47^ as well as BAL fluid^48–50^, but no study to date has directly compared sample approaches. Based on the findings of the current study, we strongly recommend whole lung tissue as a preferred sampling strategy for subsequent murine lung microbiome studies.

While BAL fluid in mice contains weak bacterial signal relative to lung tissue, in humans the opposite has been observed: human BAL specimens contain consistently stronger bacterial signal than lung tissue acquired via biopsy. This observation is consistent with anatomic and ecologic differences across species. Anatomically, human lungs are much larger than murine lungs, providing increased surface area for sampling (∼75 m^2^ vs. 0·008 m^2^) and more airspace (6 L vs. 0·001 L) to accommodate the collection of far larger volumes of BAL fluid^29,30,51^. Biopsy specimens of human lungs are typically small in volume and peripheral in anatomic location, meaning they are predominantly composed of interstitium rather than airways and alveolar space (where bacteria are more likely to be found). In contrast, use of whole lung homogenate in mice ensures capture of all bacterial DNA within the entire respiratory tract. Thus, anatomic and ecologic differences between humans and mice necessitates the use of murine-specific sampling approaches, and illustrates why a “one-size-fits-all” approach to low-biomass microbiome sampling is unlikely to work: sampling strategies will need to be tailored to their specific environmental and biologic contexts.

Numerous sources of false signal can confound detection of bacterial communities in low-biomass microbiome studies, including contamination (procedural, reagent, and sequencing) and sequencing stochasticity. Salter and colleagues elegantly demonstrated the susceptibility of low-biomass samples to reagent contamination by sequencing serial dilutions of a pure bacterial culture, where increasingly diluted specimens contained increasing abundances of taxa found in the DNA isolation reagents^1^. Other sources of contamination, such as those introduced during specimen collection (e.g. bronchoscope, surgical instruments, collection tubes) or sequencing (e.g. well-to-well contamination or index switching) may also alter the taxonomic composition of low-biomass samples^52,53^. Additionally, it has recently been demonstrated via the use of sequencing replicates that sequencing stochasticity is itself a major source of variability in microbial signal in low-biomass studies^38^. Given the numerous sources of potential false signal in low-biomass microbiome studies, we do not believe this methodological challenge can be sufficiently addressed with a simple, universal solution (e.g. a single bioinformatic “decontamination” step). Rather, as illustrated in our approach, we believe the reality of microbiologic signal must be assessed within the specific ecologic context from which it is sampled, and anchored in an understanding of microbial ecology.

Several alternate approaches to false signal in low-biomass microbiome studies have been proposed. Strategies used to detect, interpret, and in some cases, eliminate, contamination have included exclusion of taxa detected in negative controls through statistical packages^54,55^ or unbiased subtraction^7^, extraction and sequencing technical replicates^38^, calculation of abundance ratios^56^, correlation analyses^57^, hierarchical clustering^58^, and building neutral models^59^. In contrast, we propose a simple analytical approach grounded in principles of microbial ecology to discriminate true microbial signal from background-derived signal. While this approach requires the use of several complementary metrics to determine the extent of background-derived signal in each specimen type, it is relatively accessible for those conducting microbiome studies due to its dependence on open-source software, conceptual familiarity to microbial ecologists, and ease of application to other low-biomass sites. Fundamentally, this approach relies on sampling the low-biomass body site of interest and comparing the size, diversity, and taxonomic composition of the microbial community identified at that low-biomass site to all potential source communities, including background signal derived from procedural, reagent, and sequencing contamination and true microbial signal derived from contiguous body sites. This approach can thus be applied to a single specimen type to discern true bacterial signal from background-derived noise, or used to compare multiple specimen types to determine the optimal sampling method in the absence of a gold standard. Our approach does not preclude the use of complementary methods (such as those mentioned above), but rather builds a foundation rooted in thorough experimental design and microbial ecology to support further analyses.

There are several limitations to our study. We selected methods of harvesting BAL fluid and whole lung tissue which have been used by our lab and others successfully, and thus cannot directly speak to other approaches (e.g. use of lung portions or pooled BAL specimens from multiple mice). Our study only tested the use of whole lung tissue and BAL fluid for the purposes of amplicon-based sequencing, and may yield different results if other sequencing methods (e.g. metagenomic sequencing) are applied. Whole lung tissue contains much more host DNA than bacterial DNA, which can confound attempts at metagenomic analyses due to the depth of sequencing required to return reliable bacterial data^60^. Given the impossibility of performing both BAL and whole lung homogenization on the same mouse, we could not perform paired analysis on the same mice. We assumed, based on prior results^6^, that co-housed mice from the same vendor and shipment should have lung bacterial communities with similar taxonomic composition. Yet it remains possible that mouse-to-mouse variation may have confounded some comparisons. Finally, despite our efforts to thoroughly account for all possible sources of bacterial signal found in both types of lung specimens, it is possible that we have not accounted for all potential source communities, including occult sources of contamination or other body sites in contact with the lungs, such as the nasopharynx and blood.

In conclusion, we here present an ecology-based analytical approach for distinguishing true bacterial signal from background contamination in low biomass microbiome studies and provide evidence supporting the use of whole lung tissue over BAL fluid in murine lung microbiome studies. The use of our ecology-based analytic approach highlights the importance of sequencing, analyzing, and reporting ample negative controls and, to the extent possible, contiguous anatomical sites or other biological source communities to assess the reality of bacterial signal in low-biomass microbiome studies.

## Abbreviations

BAL: bronchoalveolar lavage
DNA: deoxyribonucleic acid
OTU: operational taxonomic unit
PCA: principal component analysis
PCR: polymerase chain reaction
rpm: revolutions per minute
PBS: phosphate-buffered saline
ddPCR: droplet digital PCR

## Contributors

JMB participated in study design, sample collection, data acquisition, bioinformatic and statistical analysis, data interpretation, and drafted and revised the manuscript. KJH, RAM, and NRF participated in sample collection and processing and data acquisition. GBH participated in study design, data interpretation, and manuscript revision. RPD conceived the study design and participated in data interpretation and manuscript revision. All authors read and approved this version of the manuscript.

## Data sharing

The dataset supporting the results of this article has been posted to the NIH Sequence Read Archive (accession number: PRJNA644805). The script used for mothur analysis can be found at https://github.com/piyuranjan/DicksonLabScripts/blob/master/mothurGreatLakes.sh. R code and accompanying files for the microbial community analysis and statistical tests presented in this paper can be found at https://github.com/dicksonlunglab/WholeLungvBALFluid.

## Acknowledgments

This research was supported in part by the University of Michigan Medical School Host Microbiome Initiative and by work performed by The University of Michigan Microbiome Core. Figures were created in part using BioRender.com. The authors thank Piyush Ranjan, Christopher Brown, Rishi Chanderraj, and John Erb-Downward for assistance with bioinformatic and statistical analyses and William R. Branton for his assistance in generation of pilot data.

