## Supplemental Data and Methods for "Application of an ecology-based analytic approach to discriminate signal and noise in low-biomass microbiome studies: whole lung tissue is the preferred sampling method for amplicon-based characterization of murine lung microbiota"

**Supplemental Methods**

**Mice:** Eight-week-old female C57BL/6 mice (n = 20) were purchased from Jackson Laboratories and housed under specific pathogen–free conditions. Mice were housed in five-animal cages in a common animal housing room and did not receive independent ventilation. Mice were allowed to acclimate for 1 week before harvest at 9 weeks of age. Animal experimentation was performed in compliance with the ARRIVE Guidelines[1,2].

**Tissue collection and processing:** On day of harvest, mice were randomized to either the whole lung tissue or BAL fluid groups. To account for cage effect during downstream analysis, two or three mice per five-mouse cage were randomly assigned to each sampling group, for a total of 10 mice from each of the four cages in the whole lung tissue and BAL fluid groups. Mice were sacrificed via $CO_2$ asphyxiation and organs were harvested in order of increasing biomass (tongue, whole lung tissue or BAL fluid, and cecum, respectively). All tissue samples were collected using sterile technique and instruments were rinsed with ethanol and flamed between each organ. Tongue (n=20) and cecum (n=20) samples, which were collected to serve as low- and high-biomass positive controls, respectively, were immediately snap-frozen using liquid nitrogen and stored at -80°C until DNA isolation. Both types of lung samples collected for this study were harvested and processed according to previously published protocols[3,4]. Murine whole lung tissue (n=10) was excised, placed in tubes containing 1 mL sterile water, and mechanically homogenized using a Tissue-Tearor (Biospec Products, Bartlesville, OK). The tissue homogenizer was cleaned and rinsed in ethanol and water between each tissue sample. Water control specimens from homogenization (n=2) rinsed with clean instruments, were included as procedural controls for whole lung tissue. Lung homogenate was centrifuged at 30,000 x g rpm, supernatant was removed, and the cellular pellet was snap-frozen with liquid nitrogen and stored at -80°C until DNA isolation. BAL fluid (n=10) was prepared by collecting two serial lavages of 1 mL sterile phosphate-buffered saline (PBS), yielding up to 2 mL total BAL fluid per mouse. BAL fluid was centrifuged at 30,000 x g rpm for 30 minutes, supernatant was removed, and the cellular pellet was snap-frozen with liquid nitrogen and stored at -80°C until DNA isolation. Sterile PBS (n=2) used for lavage and PBS rinses (n=4) of the syringe, and tubing collected pre- and post-lavage, were collected as procedural controls for BAL fluid.

**DNA isolation:** DNA isolation was performed with a single kit according to a modified protocol previously demonstrated to isolate bacterial DNA[5]. Briefly, genomic DNA was extracted from mouse tissue samples using a DNeasy Blood & Tissue kit (Qiagen, Hilden, Germany, catalog no. 69506) and homogenized in PowerBead Tubes (Qiagen, Hilden, Germany, catalog no. 13123-50). To detect contamination introduced by the DNA isolation kit, elution (AE) buffer (n=6) and specimen-free DNA isolations using empty bead tubes (n=6) were collected and sequenced as negative controls. Samples were processed in a randomized order to reduce false pattern formation due to reagent contamination[6].

**Bacterial DNA quantification:** Bacterial DNA in lung specimens and negative controls was quantified with a QX200 ddPCR system (Bio-Rad, Hercules, CA) according to a previously published protocol[7]. Sampling and DNA isolation controls and sterile PCR-grade water used for sample dilution as a no template control (n=4) were run alongside lung specimens. All lung specimens and negative controls were run with two technical replicates. Droplets were generated using an automated droplet generator (Bio-Rad, catalog no. 1864101). PCR amplification was performed with the Bio-Rad C1000 Touch Thermal Cycler (catalog no. 1851197). Primers were 5'- GCAGGCCTAACACATGCAAGTC-3' (63F) and 5'-CTGCTGCCTCCCGTAGGAGT-3' (355R). The cycling protocol was 1 cycle at 95°C for 5 minutes, 40 cycles at 95°C for 15 seconds and 60°C for 1 minute, 1 cycle at 4°C for 5 minutes, and 1 cycle at 90°C for 5 minutes, with all steps at a ramp rate of 2°C/second. Droplets were detected using the automated droplet reader (Bio-Rad, catalog no. 1864003), quantified using Quantasoft™ Analysis Pro (version 1.0.596), and imported to R for visualization and statistical analysis.

**16S rRNA gene sequencing:** The V4 region of the 16S rRNA gene was amplified using published primers[8] and the dual-indexing sequencing strategy developed by the laboratory of Patrick D. Schloss[9], according to the manufacturer's instructions with modifications found in the Schloss standard operating procedures (SOP)[10] as published previously[11,12]. For primary PCR, each 20 uL PCR reaction contained the following: 5 uL of a 4 uM equimolar primer set, 2 uL 10X AccuPrime PCR Buffer II (Life Technologies, catalog no. 12346094), 9.85 uL sterile PCR-grade water, 0.15 uL Accuprime High Fidelity Taq Polymerase (Life Technologies catalog no. 12346094), and 3 uL of template DNA. PCR cycling conditions were 95°C for 2 minutes, followed by 20 cycles of touchdown PCR (95°C 20 seconds, 60°C 15 seconds and decreasing 0.3 degrees each cycle, 72°C 5 minutes), then 20 cycles of standard PCR (95°C for 20 seconds, 55°C for 15 seconds, and 72°C for 5 minutes), and finished with 72°C for 10 minutes. PCR products were visualized using an E-Gel 96 with SYBR Safe DNA Gel Stain, 2% (Life Technologies catalog no. G7208-02). Samples that did not amplify during the first round of PCR were reamplified during a single round of troubleshooting using the same PCR cycling protocol and reaction composition as described above, except for the following modifications: increasing template DNA volume from 3 uL to 5 uL and decreasing the sterile PCR-grade water volume by 2 uL to yield a total reaction volume of 20 uL. After confirming successful amplification of all samples, libraries were normalized using SequalPrep Normalization Plate Kit (Life Technologies, catalog no. A10510-01) following the manufacturer's protocol for sequential elution. The concentration of the pooled samples was determined using Kapa Biosystems Library Quantification kit for Illumina platforms (Kapa Biosystems, catalog no. KK4824) and amplicon size was determined using the Agilent Bioanalyzer High Sensitivity DNA analysis kit (catalog no. 5067-4626). Libraries were prepared according to Illumina's "Preparing Libraries for Sequencing on the MiSeq" protocol for 2 nM libraries (part no. 15039740 Rev. D). The final library consisted of equimolar amounts from each of the plates normalized to the pooled plate at the lowest concentration. The final load concentration was 5 pM, spiked with PhiX at 15% to add diversity. Sequencing reagents were prepared according to the Schloss SOP and custom read 1, read 2, and index primers were added to the reagent cartridge. Amplicons were sequenced using the Illumina MiSeq platform (San Diego, CA) using a MiSeq Reagent Kit V2 (Illumina, catalog no. MS102-2003) for 500 cycles. Sterile water (n=8) and empty wells (n=28) were sequenced as negative controls and a synthetic community (n=4; ZymoBIOMICS Microbial Community DNA Standard, Zymo Research catalog no. D6306) was sequenced as a positive control. FASTQ files were generated with paired end reads and retained for further analysis.

**Adequacy of sequencing:** The full dataset obtained from the sequencing run included 5,560,120 total bacterial reads, with a mean ± SD of 46,334 ± 63,233 reads per specimen and a range of 53–287,832 reads per specimen. Except for one specimen, all tissue and control samples met the minimum requirement for number of sequencing reads and were retained for further analysis (Supplementary Figure 1). One whole lung tissue sample identified as having an insufficient number of sequencing reads (far lower than negative control samples) was excluded from further analysis. No major differences between the conclusions drawn from hypothesis tests conducted with the full sequencing dataset compared to the trimmed, quality-checked sequencing dataset were observed (Supplementary Table 1).

**Data analysis:** 16S rRNA gene sequencing data were processed using mothur (v. 1.43.0) according to the Standard Operating Procedure for MiSeq sequence data using a minimum sequence length of 250 base pairs[10,13]. A shared community file and a genus-level phylotyping file were generated using operational taxonomic units (OTUs) binned at 97% identity, using

SILVA (v. 132) for sequence alignment (silva.nr_v132.regionV4.align). OTU numbers were arbitrarily assigned in the binning process and are referred to throughout the manuscript in association with their most specified level of taxonomy (typically genus or family). OTUs were classified using the mothur implementation of the Ribosomal Database Project (RDP) classifier and RDP taxonomy training set 16 (trainset16_022016.rdp.fasta, trainset16_022016.rdp.tax), available on the mothur website[10]. After data processing with mothur, shared community (OTU) and taxonomy files were imported to R for trimming and quality checks. OTUs that composed greater than 0.1% of reads in all samples were retained in the trimmed dataset for further analysis. One sample (WVB_Lung_L3) yielded less than 100 reads and was removed from the quality-checked dataset; all other experimental and control samples were retained in the quality-checked dataset, which was used for the main analysis.

Microbial community analysis of the quality-checked dataset was performed in R[14] and relied primarily on the tidyverse (v. 1.3.0)[15], ggplot2 (v. 3.3.0)[16], vegan (v. 2.5-6)[17], and cbmbtools (v. 0.0.09025)[18] packages. For relative abundance, samples were normalized to the percent of total reads and analysis was restricted to OTUs that were present at greater than 0.1% of the sample population. No OTUs were excluded from the dataset to account for background contamination. Diversity comparisons were performed by calculating community richness rarefied to 1000 reads per sample, Shannon diversity index, and the Bray-Curtis dissimilarity index. Ordinations were performed using principal component analysis on Hellinger-transformed OTU count tables generated using Euclidean distances[19].

Overall significance was determined as appropriate by the Kruskal-Wallis test and by permutational multivariate ANOVA (PERMANOVA) with 10,000 permutations using Euclidean distances (adonis). Pairwise significance was determined as appropriate by the Wilcoxon test with the Benjamini-Hochberg correction for multiple comparisons, Tukey's HSD test, and two-sample independent Mann-Whitney U test. All statistical tests used $p=0.05$ as a threshold for significance.

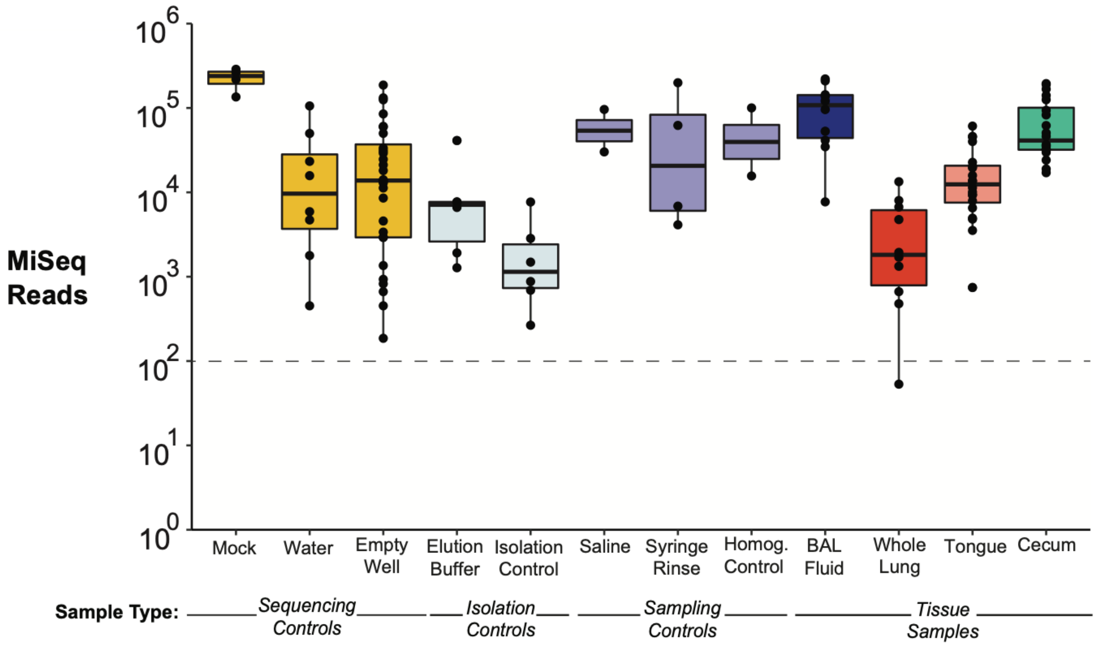


**Supplementary Figure 1: Quality 16S rRNA gene sequencing data was obtained for all sample types.** A. Sufficient numbers of reads were obtained for all sample types. Dotted line represents the number of identified reads required to be included in further analysis (minimum read count ≥ 100). One whole lung tissue specimen with 53 reads (shown below the dotted line) did not meet the minimum requirement for read count and was excluded from the main analysis. Median and IQR are shown. Individual points represent number of bacterial MiSeq reads obtained for each tissue or control specimen.

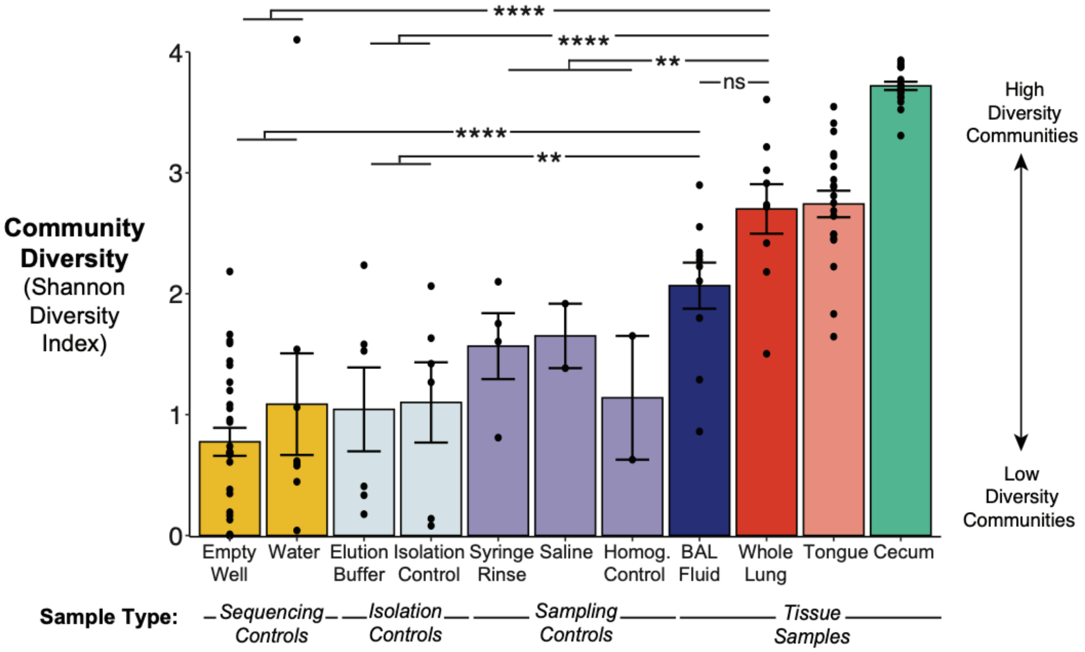


**Supplementary Figure 2: Increased alpha diversity of bacterial communities in murine whole lung tissue is driven by increased number of unique taxa relative to BAL fluid and negative controls.** Within-sample diversity of bacterial communities in whole lung tissue was comparable to that of tongue specimens and significantly greater than that of sampling, isolation, and sequencing controls. In contrast, the bacterial communities in BAL fluid were not significantly more diverse than sampling controls. Within-sample diversity of bacterial communities was quantified using the Shannon diversity index, which accounts for both richness and evenness of species diversity. When considered together with Figure 3, the difference in alpha diversity between whole lung tissue and BAL fluid appears to be driven by increased richness (number of unique OTUs) in whole lung tissue, with comparable evenness in both specimen types resulting in a non-significant difference in Shannon diversity index. Mean ± SEM and individual data points are shown. Pairwise significance was determined by comparing whole lung tissue and BAL fluid to pooled sampling, isolation, and sequencing controls (respectively, as shown) using Tukey's HSD test. Significance key: ns $p > 0.05$; * $p \leq 0.05$; ** $p \leq 0.01$; *** $p \leq 0.001$; ****$p \leq 0.0001$.

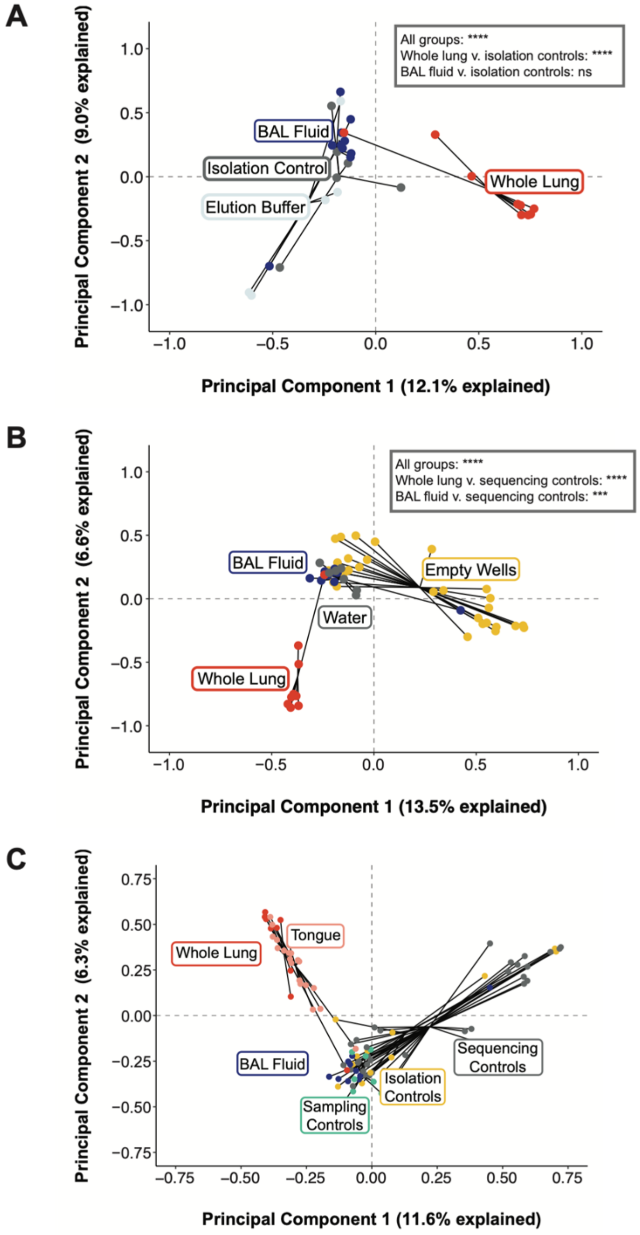


**Supplementary Figure 3: The taxonomic composition of bacterial communities in murine whole lung tissue is distinct from isolation and sequencing controls.** A. Whole lung tissue clusters separately from BAL fluid and isolation controls. B. Whole lung tissue clusters separately from BAL fluid and sequencing controls. C. Clustering of BAL fluid with negative controls and clustering of whole lung tissue with tongue specimens is evident when plotting all low-biomass specimens. For all panels, plots were generated by principal component analysis of Hellinger-transformed 16S rRNA gene sequencing data. Points represent individual specimens grouped by sample or control type. Overall significance was determined by permutational multivariate ANOVA. Pairwise significance was determined by two-sample PERMANOVA, conducted by pooling the isolation or sequencing controls, respectively, and comparing to each lung sample type. Significance key: ns $p > 0.05$; * $p \leq 0.05$; ** $p \leq 0.01$; *** $p \leq 0.001$; ****$p \leq 0.0001$.

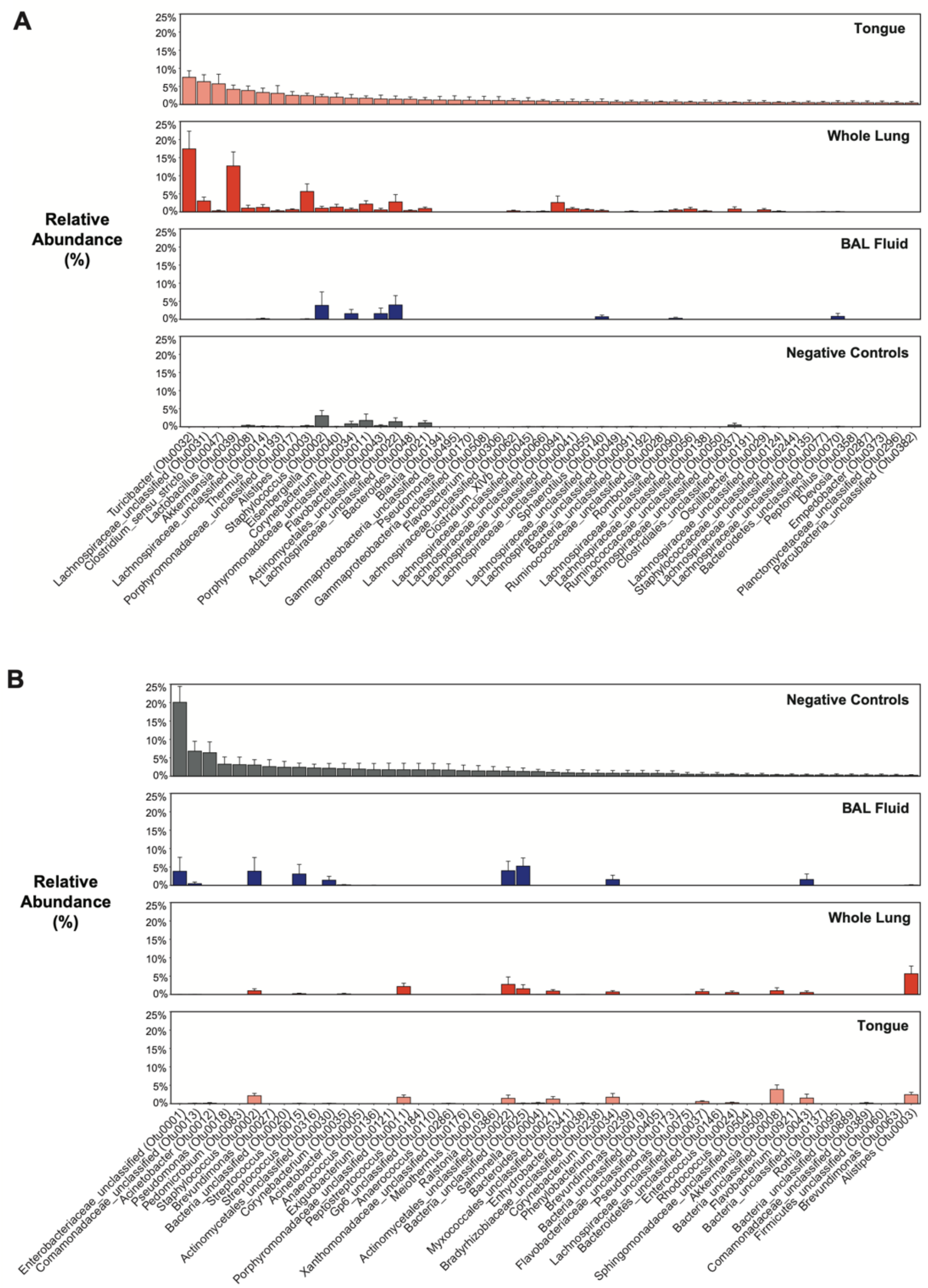


**Supplementary Figure 4: Murine whole lung tissue displays OTU-level similarity to the oral microbiome and BAL fluid displays OTU-level similarity to negative controls.** A. Whole lung tissue displays greater overlap in OTUs present in tongue specimens than does BAL fluid. B. BAL fluid displays a greater number of OTUs present in negative controls relative to whole lung tissue. Percent relative abundance of OTUs ranked by average abundance in (A) tongue specimens or (B) pooled sampling, isolation, and sequencing negative controls. Bars represent mean abundance ± SEM of the top 50 OTUs present in (A) tongue specimens or (B) pooled negative controls, found in each sample type. Labels denote genus (or most specific taxonomic level if no genus was assigned) and unique identifier for each OTU.

### Supplementary Table 1: Comparison of hypothesis test results from raw, trimmed, and final sequencing datasets

| Figure | Statistical Test | Comparison Groups | Untrimmed (Raw) Dataset | Trimmed Dataset | Trimmed & Quality-Checked (Final) Dataset |
|---|---|---|---|---|---|
| 3 | Tukey's HSD | whole lung v. BAL fluid | 0.0150895 (*) | 0.0150895 (*) | **0.0014826 (**)** |
| 3 | Tukey's HSD | whole lung v. sampling controls | 0.0087438 (**) | 0.0087438 (**) | **0.0008519 (***)** |
| 3 | Tukey's HSD | whole lung v. isolation controls | 0.0000253 (****) | 0.0000253 (****) | **0.0000011 (****)** |
| 3 | Tukey's HSD | whole lung v. sequencing controls | 0.0000000 (****) | 0.0000000 (****) | **0.0000000 (****)** |
| 3 | Tukey's HSD | BAL fluid v. sampling controls | 0.9958863 (ns) | 0.9958863 (ns) | **0.9950044 (ns)** |
| 3 | Tukey's HSD | BAL fluid v. isolation controls | 0.4262449 (ns) | 0.4262449 (ns) | **0.3746804 (ns)** |
| 3 | Tukey's HSD | BAL fluid v. sequencing controls | 0.0757056 (ns) | 0.0757056 (ns) | **0.0545862 (ns)** |
| 4 | BH-corrected pairwise Wilcoxon | whole lung v. BAL fluid | $3.8 \times 10^{-12}$ (****) | $1.3 \times 10^{-10}$ (****) | **$6.3 \times 10^{-9}$ (****)** |
| 4 | BH-corrected pairwise Wilcoxon | whole lung v. empty wells | $7.1 \times 10^{-7}$ (****) | $5.2 \times 10^{-7}$ (****) | **$7.4 \times 10^{-6}$ (****)** |
| 4 | BH-corrected pairwise Wilcoxon | BAL fluid v. empty wells | 0.29581 (ns) | 0.27300 (ns) | **0.2730 (ns)** |
| 5A | Multivariate PERMANOVA | all | 0.000099 (****) | 0.000099 (****) | **0.000099 (****)** |
| 5A | Two-sample PERMANOVA | whole lung v. negative controls | 0.0002 (***) | 0.0002 (***) | **0.0004 (***)** |
| 5A | Two-sample PERMANOVA | whole lung v. BAL fluid | 0.000099 (****) | 0.000099 (****) | **0.000099 (****)** |
| 5A | Two-sample PERMANOVA | BAL fluid v. negative controls | 0.4382 (ns) | 0.464 (ns) | **0.463 (ns)** |
| 5B | Multivariate PERMANOVA | all | 0.000099 (****) | 0.000099 (****) | **0.000099 (****)** |
| 5B | Two-sample PERMANOVA | whole lung vs. tongue | 0.009199 (**) | 0.009199 (**) | **0.0142 (*)** |
| 5B | Two-sample PERMANOVA | BAL fluid vs. tongue | 0.000099 (****) | 0.000099 (****) | **0.000099 (****)** |
| 5C | Mann-Whitney U test | whole lung vs. BAL fluid | 0.0002057 (***) | 0.0002057 (***) | **0.0004114 (***)** |
| S2 | Tukey's HSD | whole lung v. BAL fluid | 0.3728207 (ns) | 0.3602822 (ns) | **0.3098538 (ns)** |
| S2 | Tukey's HSD | whole lung v. sampling controls | 0.0071118 (**) | 0.0077170 (**) | **0.0067338 (**)** |

| Figure | Statistical Test | Comparison Groups | Untrimmed (Raw) Dataset | Trimmed Dataset | Trimmed & Quality-Checked (Final) Dataset |
|---|---|---|---|---|---|
| S2 | Tukey's HSD | whole lung v. isolation controls | 0.0000143 (****) | 0.0000190 (****) | **0.0000205 (****)** |
| S2 | Tukey's HSD | whole lung v. sequencing controls | 0.0000000 (****) | 0.0000000 (****) | **0.0000000 (****)** |
| S2 | Tukey's HSD | BAL fluid v. sampling controls | 0.3911169 (ns) | 0.4191854 (ns) | **0.4234102 (ns)** |
| S2 | Tukey's HSD | BAL fluid v. isolation controls | 0.0106434 (*) | 0.0141500 (*) | **0.0147639 (*)** |
| S2 | Tukey's HSD | BAL fluid v. sequencing controls | 0.0000534 (****) | 0.0000800 (****) | **0.0000876 (****)** |
| S3A | Multivariate PERMANOVA | all | 0.000099 (****) | 0.000099 (****) | **0.000099 (****)** |
| S3A | Two-sample PERMANOVA | whole lung vs. isolation controls | 0.000099 (****) | 0.000099 (****) | **0.000099 (****)** |
| S3A | Two-sample PERMANOVA | BAL fluid vs. isolation controls | 0.05969 (ns) | 0.07029 (ns) | **0.07149 (ns)** |
| S3B | Multivariate PERMANOVA | all | 0.000099 (****) | 0.000099 (****) | **0.000099 (****)** |
| S3B | Two-sample PERMANOVA | whole lung vs. sequencing controls | 0.000099 (****) | 0.000099 (****) | **0.000099 (****)** |
| S3B | Two-sample PERMANOVA | BAL fluid vs. sequencing controls | 0.0004 (***) | 0.0004 (***) | **0.0004 (***)** |

**Supplementary Table 1: Hypothesis test results were consistent across datasets at each stage of quality filtering.** All values indicate the p-value from the specified comparison using the datasets indicated. The untrimmed (raw) dataset refers to the sequencing data as processed by mothur before exclusion of low-frequency OTUs and samples with insufficient reads. The trimmed dataset refers to the raw dataset after exclusion of low-frequency OTUs, but without exclusion of any samples; OTUs that comprised greater than 0.1% of reads for one or more samples were retained in the trimmed dataset. The trimmed and quality-checked (final) dataset refers to the trimmed dataset after exclusion of samples with insufficient numbers of reads; samples ≥ 100 reads were retained in the quality-checked dataset. All experimental and controls samples except one whole lung tissue specimen were retained in the quality-checked datasets (see Supplementary Figure 1). Bolded p-values indicate comparisons included in the referenced figure.
